# Sensory and perceptual decisional processes underlying the perception of reverberant auditory environments

**DOI:** 10.1101/2024.03.12.584683

**Authors:** Haydée G García-Lázaro, Santani Teng

**Author notes:** **Address for correspondence:** The Smith-Kettlewell Eye Research Institute 2318 Fillmore St., San Francisco, CA, 94115.

## Abstract

Reverberation, a ubiquitous feature of real-world acoustic environments, exhibits statistical regularities that human listeners leverage to self-orient, facilitate auditory perception, and understand their environment. Despite the extensive research on sound source representation in the auditory system, it remains unclear how the brain represents real-world reverberant environments. Here, we characterized the neural response to reverberation of varying realism by applying multivariate pattern analysis to electroencephalographic (EEG) brain signals. Human listeners (12 male and 8 female) heard speech samples convolved with real-world and synthetic reverberant impulse responses and judged whether the speech samples were in a “real” or “fake” environment, focusing on the reverberant background rather than the properties of speech itself. Participants distinguished real from synthetic reverberation with ∼75% accuracy; EEG decoding reveals a multistage decoding time course, with dissociable components early in the stimulus presentation and later in the peri-offset stage. The early component predominantly occurred in temporal electrode clusters, while the later component was prominent in centro-parietal clusters. These findings suggest distinct neural stages in perceiving natural acoustic environments, likely reflecting sensory encoding and higher-level perceptual decision-making processes. Overall, our findings provide evidence that reverberation, rather than being largely suppressed as a noise-like signal, carries relevant environmental information and gains representation along the auditory system. This understanding also offers various applications; it provides insights for including reverberation as a cue to aid navigation for blind and visually impaired people. It also helps to enhance realism perception in immersive virtual reality settings, gaming, music, and film production.

**SIGNIFICANCE:** In real-world environments, multiple acoustic signals coexist, typically reflecting off innumerable surrounding surfaces as reverberation. While reverberation is a rich environmental cue and a ubiquitous feature in acoustic spaces, we do not fully understand how our brains process a signal usually treated as a distortion to be ignored. When asking human participants to make perceptual judgments about reverberant sounds during EEG recordings, we identified distinct, sequential stages of neural processing. The perception of acoustic realism first involves encoding low-level reverberation acoustic features and their subsequent integration into a coherent environment representation. This knowledge provides insights for enhancing realism in immersive virtual reality, music, and film production, and using reverberation to guide navigation for blind and visually impaired people.

## INTRODUCTION

In real-world acoustic environments, listeners receive as inputs combined signals of sound sources (e.g., speech, music, tones, etc.) and their reflections from surrounding surfaces. These reflections are attenuated, time-delayed, and additively aggregated as reverberation, a cue that listeners leverage to self-orient (Flanagin et al., 2017; Kolarik, Cirstea, et al., 2013; Kolarik, Pardhan, et al., 2013), optimize perception (Bidelman et al., 2018; Francl & McDermott, 2022; Slama & Delgutte, 2015)2015), and make inferences about the environment (Kolarik et al., 2021; Papayiannis et al., 2020; Peters et al., 2012; Shabtai et al., 2010; Teng et al., 2017; Traer et al., 2021).

Real-world reverberation exhibits consistent statistical regularities: it is dynamic, bound to the sound source envelope, and decays exponentially with a frequency-dependent profile. In contrast, background noise with deviant decay or spectral profiles is more difficult to segregate and can be distinguished as synthetic (Traer & McDermott, 2016). This suggests that the auditory system carries a relatively low-dimensional but finely tuned representation of natural environmental acoustics, distinct from source sounds, other acoustic backgrounds, and noise (Francl & McDermott, 2022; Kell & McDermott, 2019). However, perceiving sounds as authentic requires more than replicating the low-level regularities of natural signals. For example, observers still do not always perceive physically faithful synthetic signals as completely authentic, even though brain responses accurately classify them (Moshel et al., 2022). This suggests that other constraints and contextual information drive auditory judgments in addition to physically realistic acoustics. Thus, perceiving natural acoustic environments likely involves both low-level sensory encoding and higher-level cognitive processes.

The neural representations of real-world acoustic environments remain largely unexplored. Most research has focused on the robustness of neural representations of sound sources against background signals (Fujihira et al., 2017; Mesgarani et al., 2014; Puvvada et al., 2017; Sayles & Winter, 2008). These representations become more invariant to background noise and reverberations along the auditory system hierarchy (Fuglsang et al., 2017; Kell & McDermott, 2019; Khalighinejad et al., 2019; Lowe et al., 2022; Mesgarani et al., 2014; Rabinowitz et al., 2013; Slama & Delgutte, 2015). Notably, brain responses to reverberant sounds—the combination of sound source and reverberation—encode both the source and the size of the space or the environment separately, even when the reverberant signal is not relevant to the task at hand (Flanagin et al., 2017; Puvvada et al., 2017; Teng et al., 2017). Cortical responses to slightly delayed echoic speech seem to represent the anechoic speech envelope, presumably to aid intelligibility (Gao et al., 2024). This suggests that sound sources and reverberation may be segregated and represented by different neural codes along the auditory pathway (Puvvada et al., 2017). The mechanisms of segregation and the temporal and spatial scales at which these operations occur are still under study (Devore et al., 2009; Ivanov et al., 2021; Rabinowitz et al., 2013). The direct link between the neural representations of acoustic features and perception of reverberation remains unknown, likely because almost no study has focused on reverberation as a target signal, but rather as a noise-type signal accompanying other sound sources.

Here, we characterized neural responses to reverberant acoustic environments of varying realism. We recorded electroencephalographic (EEG) signals while participants performed an auditory task with physically realistic or unrealistic reverberations as part of the stimuli.

Participants determined whether speech samples were in a real or synthetic environment, i.e., judging the properties of the reverberant background rather than those of the sound source.

We expected neural responses to encode acoustic statistics and listeners’ behavioral judgments distinguishing real and synthetic ("fake") reverberation. We further expected neural response patterns to reverberation to behave dynamically, with functionally distinct components reflecting the underlying operations of the perceived realism judgments. Our results show that neural responses track the spectrotemporal regularities of reverberant signals in sequential and dissociable stages, likely reflecting sensory encoding and higher perceptual decision-making processes.

## METHODS

### Participants

Twenty healthy young volunteers (mean age = 32.3 years, SD = 5.7 years, 12 male) participated in this experiment. All participants had normal or corrected to normal vision, reported normal hearing, and gave informed consent in accordance with a protocol approved by the Smith-Kettlewell Eye Research Institute Institutional Review Board.

### Stimuli

Stimuli were 2-second extracts of compound sounds created by convolving sound sources (spoken sentences) with reverberant impulse responses (IRs). The speech samples were taken from the TIMIT Acoustic-Phonetic Continuous Speech Corpus (John S. Garofolo, Lori F. Lamel, William M. Fisher, Jonathan G. Fiscus, David S. Pallett, Nancy L. Dahlgren, Victor Zue, 1993), equally balanced between male and female speakers and unique to each of the 600 stimuli. The IRs comprised the 30 most reverberant signals from a collection of real-world recordings by Traer & McDermott (Traer & McDermott, 2016) (Mean RT_60_ 1.0686 sec, min = 0.8587 sec, max = 1.789 sec, with no difference in decodability between them, Figure 1 in supplementary material). Additionally, we generated five synthetic IRs for each real-world (*Real*) IR that reproduced or altered various temporal or spectral statistics of the *Real* IRs’ reverberant tails (omitting early reflections), adapting the code and procedures detailed in Traer & McDermott (Traer & McDermott, 2016). Briefly, Gaussian noise was filtered into 32 spectral subbands using simulated cochlear filters, and an appropriate decay envelope was imposed on each subband. Temporal variants were created by reversing the original IR profile (*Time-Reversed*) or imposing a linear rather than exponential decay envelope onto each subband (*Linear Decay*). Spectral variants were created by preserving exponential decay but manipulating the spectral dependence profile so that middle frequencies decayed faster than lows and highs (*Inverted Spectral Dependence*) or flattening the profile so that all frequency subbands decayed equally (*Flat Spectral Dependence*). A fifth synthetic condition, *Ecological*, preserved the spectral and temporal statistics of the Real-world IR. Thus, each of the 30 real-world IRs was convolved with 5 unique speech samples and each of the 5 synthetic variants with a unique sample, for a total of 600 unique convolved sounds, half with *Real* and half with *Synthetic* reverberation (see Figure 1A-B). Finally, to equate overall stimulus dimensions between trials, we extracted 2-second segments from each convolved sound, yoked to the local amplitude peak within the first 1000 ms, RMS-equated, and ramped with 5-ms on- and offsets. Thus, judging the stimulus category required perceptual extraction of the reverberation itself, as it was not predicted by overall stimulus duration, intensity, speech sample, or speaker identity.

**Figure 1.**
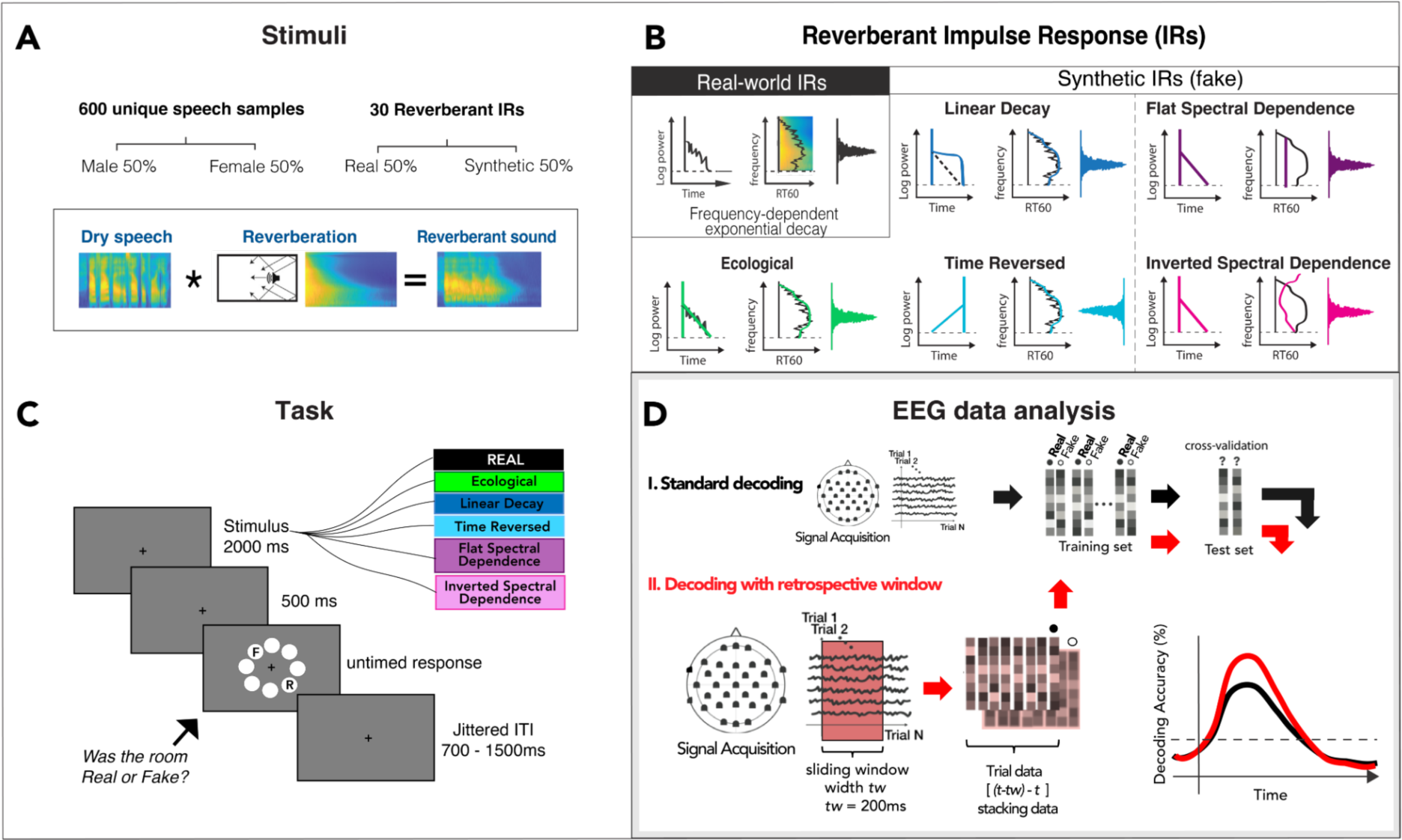
Experimental design. A: Stimuli were created by convolving 600 unique dry speech samples (50% male and 50% female voices) with one of 30 reverberant impulse responses (IRs). B: Reverberant IRs comprised Real-world reverberations and their Synthetic (“Fake”) variants (*Ecological, Linear Decay, Time-Reversed, Flat- and Inverted-Spectral Dependence*). C: The task consisted of listening to a reverberant sound lasting 2000 ms, followed by a 500 ms blank before an untimed response display appeared. Subjects judged whether the reverberation was Real or Synthetic (“Fake”) by clicking on the “R” or F,” whose locations varied pseudo-randomly trial by trial to prevent motor response preparation before the response cue. Responses were followed by a jittered interstimulus interval (ITI) varying between 700 and 1500 ms. D: EEG data analysis pipeline: Multivariate Pattern Analysis using retrospective sliding window.

### Task and Experimental Procedure

The experiment was conducted in a darkened, sound-damped testing booth (Vocalbooth, Bend, OR). Participants sat 60 cm from a 27" display (Asus ROG Swift PG278QR, Asus, Taipei, Taiwan), wearing the EEG cap (see below) and tubal-insert earphones (Etymotic ER-3A). Stimuli were monaural and presented diotically. To improve the temporal precision of auditory stimulus presentation, we used the MOTU UltraLite Mk4 Audio Control (MOTU Inc., Cambridge, MA) as an interface between the stimulation computer and the earphones. Sound intensity was jittered slightly across trials.

Participants performed a 1IFC categorical judgment task, reporting whether the reverberant background of each stimulus was naturally recorded ("Real") or synthesized ("Fake"). Each trial consisted of a 2-second reverberant stimulus, 500 ms post-offset, and an untimed response window. Prior to the response cue, the display consisted only of a central fixation cross on a gray background. The response window comprised a circular array of 8 light gray circles, positioned at 45° intervals with a radius of ∼3.3° from the display center. The letters "R" and "F," corresponding to the response choices, were displayed on two diametrically opposed circles along a randomized axis on each trial (Figure 1C). Virtual response "buttons" were thus a consistent angular distance away from each other and the display center, but at a location unknown to the participant until the response window began. Responses were collected via mouse click. Thus, each trial’s stimulus condition and reported percept were decoupled from response-related movements or motor preparation. Responses were followed by a jittered interstimulus interval of 700–1500 ms before the next trial. Trials were presented in 10 blocks of 60 trials each, randomizing IRs across blocks and counterbalancing Real and synthetic IRs within a block. Total experiment time was 75–90 minutes, including rests between blocks. Stimulus presentation was programmed in Psychtoolbox-3 (Pelli, 1997) running in Matlab 2018b (The Mathworks, Natick, MA).

### EEG data acquisition and preprocessing

Continuous EEG was recorded using a Brain Products actiCHamp Plus recording system (Brain Products GmbH, Gilching, Germany) with 32 to 64 channels arranged in a modified 10-20 configuration on the caps (Easycap GmbH, Herrsching, Germany). The *Fz* channel was used as the reference during the recording. EEG signal was band-pass filtered online from 0.01 to 500 Hz and digitized at 1000 Hz. The continuous EEG signal was preprocessed offline using Brainstorm software (Tadel et al., 2011) and customized scripts using Matlab functions for downsampling and filtering the neural signal. Raw data were re-referenced to the common average of all electrodes and segmented into epochs of -400 ms to 2500 ms relative to stimulus onset. Epochs were baseline corrected and downsampled by averaging across non-overlapping 10-ms windows (Guggenmos et al., 2018), and low-pass filtered at 30 Hz. Depending on the specific analysis, trials were labeled and variously subdivided by stimulus condition (*Real*, *Fake*), perceptual report (*perceived Real, perceived Fake*), behavioral status (*Correct, Incorrect*), or specific IR variant (*Ecological, Linear Decay, Time-Reversed, Flat Spectral, Inverted Spectral*).

### EEG Multivariate Pattern Analysis (MVPA)

We used linear support vector machine (SVM) classifiers to decode neural response patterns at each time point of the preprocessed epoch using a multivariate pattern analysis (MVPA) approach. We applied a retrospective sliding window in which the classifier for time point t was trained with preprocessed and subsampled sensor-wise data in the interval [t-20, t]. This differs from similar analyses that put the decoding in the middle of the window (Schubert et al., 2020, 2021).

Pattern vectors within the window were stacked to form a composite vector (e.g., 21 samples of 63-channel data formed a length of 1323 vectors), which was then subjected to decoding analysis (Figure 1D). This method increases the signal-to-noise ratio (SNR) and captures the temporal integration of the dynamic and non-stationary properties of reverberation. The resultant decoding time course thus began at -200 ms relative to stimulus onset. Decoding was conducted using custom Matlab scripts adapting functions from Brainstorm’s MVPA package (Tadel et al., 2011) and libsvm (Chang & Lin, 2011). We used 10-fold leave-one-out cross-validation, in which trials from each class were randomly assigned to 10 subsets and subaverages (Guggenmos et al., 2018). This procedure was repeated with 100 permutations of subaverage sets; the final decoding accuracy for *t* represents the average across permutations.

### Temporal Generalization

To further investigate the temporal dynamics of the EEG response, we generalized the 1-d decoding analysis by testing the classifiers trained at each time point at all other time points within the epoch. Temporal generalization estimates the stability or transience of neural representations by revealing how long a model trained at a given time successfully decodes neural data at other time points (King & Dehaene, 2014). In the resulting two-dimensional temporal generalization matrix (TGM), the x- and y-axes index the classifiers’ testing and training time points. The diagonal of the matrix, in which *t*_Train_ = *t*_Test_, is equivalent to the 1-d decoding curve.

### Sensor-space decoding analysis

To explore the spatiotemporal distribution of the decoding analysis, we recomputed TGMs on two hypothesis-driven subsets of electrodes. We created local clusters of non-overlapping electrodes by selecting the closest electrodes around a defined centroid electrode (T7, T8, and Pz) based on a 2D projection of the sensor’s positions using the function *ft_prepare_neighbors* with the *method “distance”* from the Fieldtrip Toolbox. (Oostenveld et al., 2011) In particular, we focused on decoding results from a cluster comprising temporal sensor positions pooled across hemispheres (FC5, FC6, FT9, FT10, C3, C4, CP5, CP6, T7, T8, TP9, and TP10) and a cluster comprising 6 centroparietal sensors (Cz, Pz, P3, P4, CP1, and CP2). In this way, we obtained 2 decoding TGMs per subject, allowing us to compare temporal and representational dynamics of the neural response across broad anatomical regions (Fyshe, 2020; Oosterhof et al., 2016). Statistical analyses were performed on single-subject data and plotted across the group averaged signals.

### Brain-Behavior Correlation

We correlated performance with condition-specific decoding accuracy to further investigate the link between participants’ neural and behavioral responses. The *Real* versus *Fake* linear SVM classification analysis was carried out separately for *Real trials vs. Ecological, Linear Decay, Time-Reversed, Flat Spectral, and Inverted Spectral* conditions. We computed a nonparametric (Spearman’s rho) correlation between each participant’s behavioral and decoding accuracy, generating a time course of the extent to which the EEG response was related to participants’ judgments.

### Statistical testing

To assess the statistical significance of the EEG decoding time courses across subjects, we used t-tests against the null hypothesis of chance level (50%). We used non-parametric permutation-based cluster-size inference to control for multiple comparison error rate inflation. The cluster threshold was set to α = 0.05 (right-tail) with 1000 permutations to create an empirical null hypothesis distribution. The significance probabilities and critical values of the permutation distribution were estimated using a Monte Carlo simulation (Maris & Oostenveld, 2007).

## RESULTS

### Natural environmental acoustics are perceptually accessible

Participants correctly classified stimuli with an overall accuracy of 75.7% (*Real*: 81.95%; *Fake*: 69.43%), as shown in Figure 2A. Wilcoxon signed-rank tests revealed that both *Real* and *Fake* classification accuracy was greater than chance (Z = 3.88, p < 0.001; Z = 3.24, p < 0.01); *Real* trials were more accurately categorized than *Fake* trials (Z = 1.99, p < 0.05). Among *Fake* IRs, performance varied from chance level to near-ceiling (Figure 2B); *Linear Decay* (Z = 3.12, p< 0.01), *Time-Reversed* (Z = 3.99, p< 0.001), and *Inverted Spectral* (Z = 2.15, p< 0.05) conditions elicited above-chance performance, while *Ecological* (Z = 1.04, p = 0.29) and *Flat Spectral* (Z = 1.6, p = 0.11) conditions accuracy were at chance. Combining both temporal conditions (*Linear Decay* and *Time Reversed*) and both spectral conditions (*Inverted Spectral* and *Flat Spectral*), we observed that listeners more accurately detected temporal than spectral variants (Z = 3.92, p< 0.0001, Figure S1, supplementary materials). Response times did not differ between the conditions (mean = 1.538 s ± s.e.m. = 0.159, across subjects). Overall, this pattern is broadly consistent with previous and ongoing studies of reverberant perception (Garcia-Lazaro et al., 2021; Traer & McDermott, 2016; Wong-Kee-You, A. M. B., Alwis, Y., & Teng, S., 2021).

**Figure 2.**
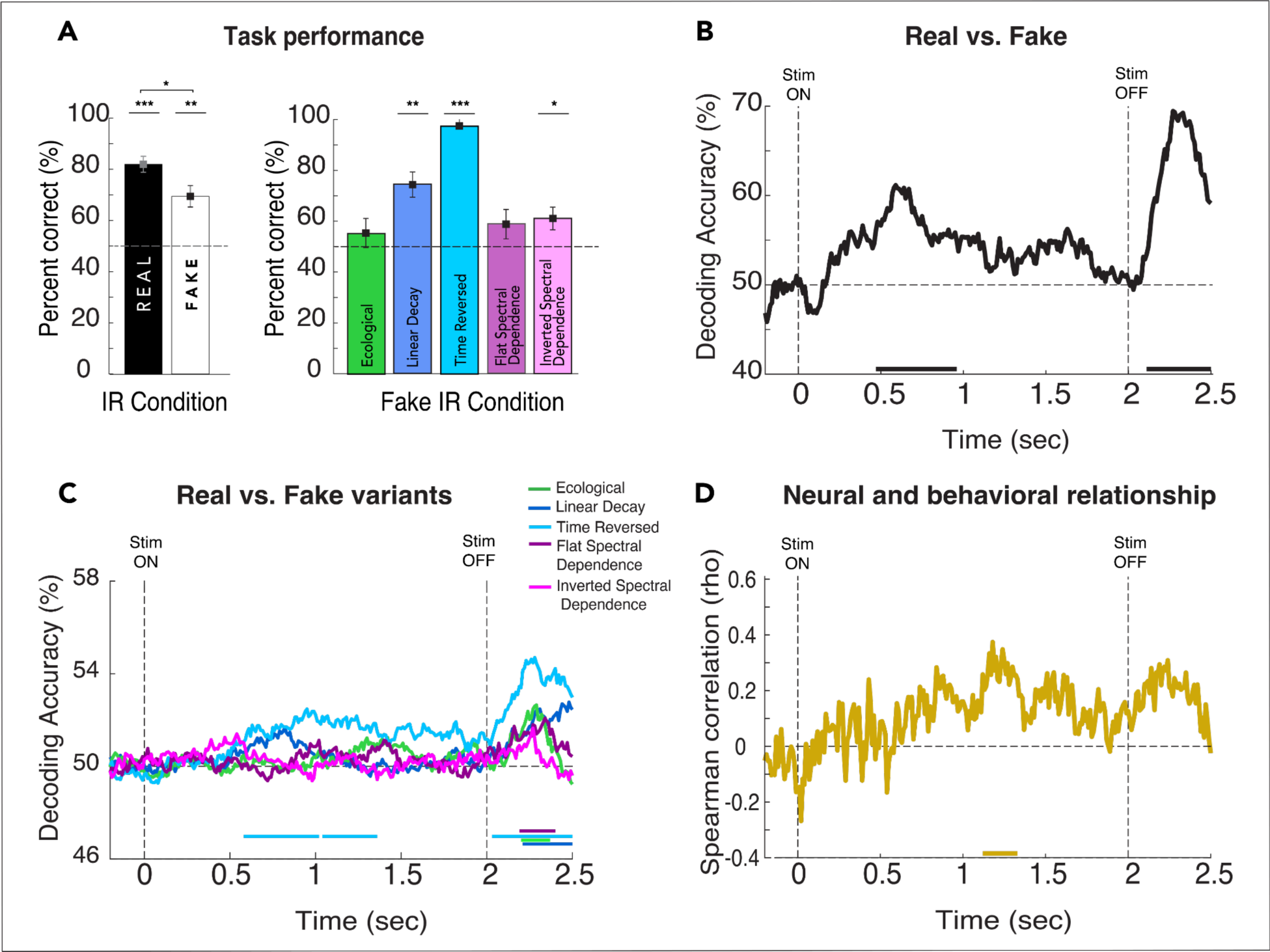
Behavioral and neural signatures of reverberant perception A: Group-level performance (mean accuracy ± s.e.m. across participants) for *Real* and *Fake* trials overall and broken down by synthetic variants B: *Real* vs. *Fake* average decoding time courses across subjects; C: Decoding time courses of *Real* vs. *Fake* variants averaged across subjects. The vertical dashed lines at zero and 2 seconds indicate stimulus onset and offset, respectively; the horizontal dashed line indicates decoding percentage at the chance (50%); and the horizontal colored bars in the x-axis indicate significance. D: Time course of brain-behavior correlation (Spearman correlation, rho) relating pairwise decoding of *Real* and *Fake* variants (panel C) to each participant’s behavioral performance (panel A). For all statistics, N = 20; *t-test* against 50%; cluster-definition threshold, *p* < 0.05; 1000 permutations. \**p* < 0.05, \*\**p* < 0.01, \*\*\**p* < 0.001.

### Neural responses track reverberant acoustics in two phases

#### EEG decoding time courses

We used linear SVM classification to compare how well *Real* and *Fake* trials could be distinguished. Figure 2B depicts the decoding time course with two reliable decodability intervals: 470–960 ms and 2110–2500 ms. The first decoding peak occurs while the stimulus is being played, while the second one peaks after the offset of the stimulus but before the response display appears. This dynamic pattern suggests that a time-evolving neural process with two critical information-processing time windows underpins reverberation perception.

To examine the neural response to each *Fake* variant relative to *Real* IRs in more detail, we applied linear SVM pairwise classification to all *Rea*l versus each *Fake* variant individually. Figure 2C shows the decoding time course for each pairwise comparison: *Real* vs. *Ecological* (green) reached significance from 2200 to 2370 ms; *Real* vs. Linear Decay (blue) reached significance from 2100 ms to 2500 ms; *Real* vs. *Time-Reversed* (light blue) reached significance from 580 ms to 1360 ms and 2030 to 2500 ms; *Real* vs. *Flat Spectral Dependence* (violet) became significant from 2190 to 2400 ms and *Real* vs. *Inverted Spectral Dependence* (pink) did not reach significance at any time points. Note that fewer trials for these condition-wise analyses may have decreased the available statistical power (⅕ of *Fake* variants). One may wonder if the *Time-Reversed* variant, which has a higher peak during the early phase of the neural response, is the primarily responsible for the early decoding peak. To rule out this possibility, we applied SVM classification to all *Real* trials and all *Fake* variants except *Time-Reversed.* As seen in Figure S2 in the supplementary material, the decoding accuracy time course for all *Fake* variants except *Time-Reversed* (blue) has similar morphology and reached significance from 530 to 780 ms and 2140 to 2500 ms, similarly as it does when comparing all *Real* versus all *Fake* variants reported in Figure 2B. The analysis is repeated, excluding each of the other synthetic variants with similar results (Supplementary Figure S2). Finally, Figure 2D shows the correlation between the condition-wise decoding and behavioral accuracy for each participant, averaged over participants of the stimulus and overall performance. Significance is reached from 1200 ms to 1350 ms after stimulus onset.

Overall, brain responses track reverberant acoustics at two distinct stages of the trial: during stimulus presentation and after stimulus offset, suggesting distinct underlying neural computations.

### Peri-offset decoding does not reflect motor response preparation or sound offset response

We have shown that the neural response to reverberant sounds is modulated in a condition-specific way during two critical time windows in the trial. To further investigate the nature of the second decoding peak, which occurs near stimulus offset, we asked whether it reflects aspects of the trial, such as a generic offset response, incidental acoustic features, motor response preparation, or mapping response location. In the first case, this response would not be unique to the *Real* vs. *Fake* comparison but would appear in other pairwise decoding results. To test this possibility and as a control against spurious features driving decoding results, we ran two comparisons. First, we decoded neural signals labeled by IR type (room type) independently of whether they were *Real* or *Fake* (Supplementary Figure S3A). Then, as a second control, we decoded neural signals according to speaker gender: male versus female, a stimulus feature orthogonal to IRs in the stimuli (Supplementary Figure S3B). Despite all trials having the same sound offset at 2 seconds, neither the room type nor speaker gender decoding analysis revealed any significant decodability.

Next, although we designed the task to decouple perceptual processing from motor preparation and mapping response location (e.g., mouse pointer movement), we tested this design empirically by repeating the decoding analysis with each trial labeled by its physical response location (out of 8 possible locations on the response array). Supplementary Figure S3C shows that decodability by response location remains at chance throughout the entire epoch of interest (-200 to 2500 ms), becoming significant only after 2500 ms when the response display actually appears.

Taken together, these analyses demonstrate that the second decoding peak reflects the experimentally manipulated reverberation signals rather than motor-related activity or incidental attributes of the stimulus.

### Physical and perceived reverberant realism are differently represented in the brain

We showed that the neural response reliably decoded *Real* versus *Fake* IR stimulus conditions. However, listeners also judged every sound, meaning that each trial had both a veridical label reflecting stimulus acoustics and a behavioral report reflecting the observer’s percept. The behavioral performance results in Figure 2 indicate that these labels differed by about 25%. Therefore, we next asked whether and how the neural response would differ as an index of observer percepts. To this end, we relabeled trials by behavioral reports (*perceived Real* versus *perceived Fake*) regardless of their veridical condition and applied the same decoding analysis described above.

Figure 3A plots the *perceived Real* vs. *Fake* decoding curve, compared against the standard physical decoding time course in black; it was significantly above chance from 1770 ms to 2500 ms, in contrast to the 470–960 ms and 2110–2500 ms regimes for physical stimulus conditions reported in Figure 2B. The shaded area indicates that decoding accuracy was significantly higher for physical stimuli than perceptual reports, from 2200 ms to 2450 ms. Overall, these results suggest that the late phase of the neural response is particularly sensitive to the subjective representation of environmental acoustics.

**Figure 3.**
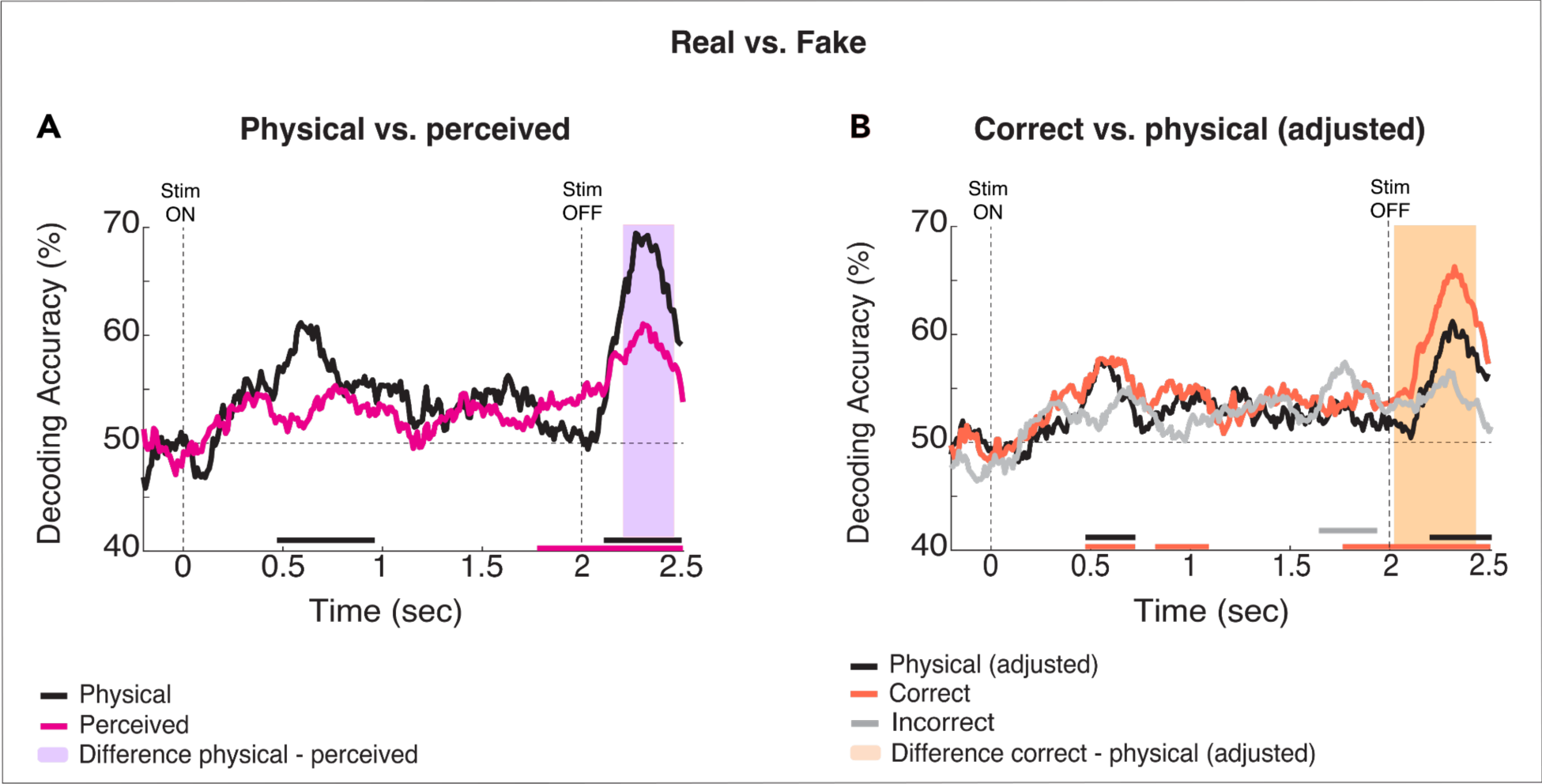
Neural signatures of reverberation perception A: Decoding accuracy time courses for *Real* versus *Fake* trials labeled by *physical* (black line) and *perceived* (pink line) categories. The vertical dashed lines at 0 and 2 sec indicate stimulus onset and offset; the horizontal dashed line indicates chance (50%), and color-coded bars in the x-axis indicate significance. The violet area denotes the time when the *physical* and *perceived* decoding differed. B. *Real* vs. *Fake* decoding accuracy for *Correct* (orange) and *Incorrect* (gray) trials, compared against *Physical* (black) adjusted to match trial count for *Correct* condition. For all statistics, N = 20; *t-test* against 50%, cluster-definition threshold, *p* < 0.05; 1000 permutations. For incorrect response trials, N = 18 because 2 subjects did not have sufficient *Real* incorrect trials.

### Correctly judged trials are more accurately decodable

We have shown that decoding time courses are sensitive to whether trials are labeled by their physical vs. reported categories. To further explore the relationship between perceived and physical reverberant acoustics, we analyzed brain responses for correctly and incorrectly judged trials, i.e., when the physical and reported categories matched and when they differed. Previous research has shown that neural signals are denoised, and neural representations of targets (IRs here) are tuned and sharpened when successfully discriminated (Alain & Arnott, 2000; Cantisani et al., 2019; King et al., 2016; Treder et al., 2014; van Bergen et al., 2015). For each subject, we first filtered trials for correct responses and then decoded them as described above. Next, we repeated the standard stimulus decoding, subsampling from all trials per subject and condition to match the number of correct trials. Figure 3B shows time courses for decoding accuracy, one in orange for *correct response* trials and one in black for *physical (adjusted) stimulus* types. Decoding accuracy for *correct response* trials was significant from 470 to 720 ms, 820 to 1090 ms, and 1760 to 2500 ms, whereas, for *physical (adjusted) stimuli*, decoding accuracy was significant from 470 to 720 ms and 2190 ms to 2500 ms. The orange shaded area in Figure 3B represents the time when the curves of *Correct* response and *physical (adjusted)* differed statistically, from 2040 ms to 2430 ms.

As expected, the number of *Incorrect* response trials was lower than the number of *Correct* ones (average across subjects: 75% *Correct* trials; 25% *Incorrect* trials; Figure 2A for reference). For this analysis, two subjects were excluded due to insufficient *Real Incorrect* response trials. Decoding accuracy for *Incorrect* trials is shown in Figure 3B. Decoding accuracy became significant between 1600 and 1890 milliseconds. Due to the large difference in the number of trials between *Correct* and *Incorrect* responses, no direct statistical comparison between these conditions is reported here.

### Reverberation perception involves temporally and functionally distinct stages of processing

The decoding time courses shown above suggest that the neural representations of reverberation evolve over distinct, sequential time windows. However, we have yet to determine whether these two regimes of significance reflect two instances of the same neural computation (e.g., a re-entrant sensory representation) or two distinct stages of the reverberant perceptual judgment. To do that, we cross-classified the EEG signal across time points to elucidate these dynamics by generating temporal generalization matrices of *Real* vs. *Fake* classifications. As described earlier, TGMs enable us to examine how transient or persistent neural representations are. The TGM reveals this information by displaying how long a model trained at one time can successfully decode neural data at other time points (King & Dehaene, 2014). Thus, we reasoned that if the two decoding peaks found in the decoding time course correspond to the same neural operation, the TGM will exhibit a pattern in which classifiers from earlier times can reliably decode brain signals from later times (generalization pattern). In contrast, if two decoding peaks are supported by two distinct operations with different dynamics, the TGM will not generalize over time; rather, it will exhibit a pattern in which classifiers from earlier times fail to classify brain signals from later times reliably (King & Dehaene, 2014).

Figure 4 A-C shows TGMs for *physical* Real vs Fake trials, *perceived Real* vs. perceived *Fake*, and *Correct Real* vs. *Correct Fake,* respectively. The TGM plots all show two regimes of significant decoding accuracy, highlighted in white: from 250–1250 ms and 1500–2500 ms (*physical*, Fig. 4A); ∼70–1500 ms and ∼1500–2500 ms (*perceived*, Fig. 4B); and starting at ∼180 ms, broadening slightly after 1500 ms until the end of the epoch (*Correct*, Fig. 4C). TGM for incorrect response trials did not achieve significance at any time during the epoch. These brain patterns correspond to two sequential windows of independent information processing, as evidenced by their non-generalized pattern across time (Cichy & Teng, 2017; Fyshe, 2020; King & Dehaene, 2014) and indicate that the perception of natural reverberant environments is underpinned by distinct neural operations.

**Figure 4.**
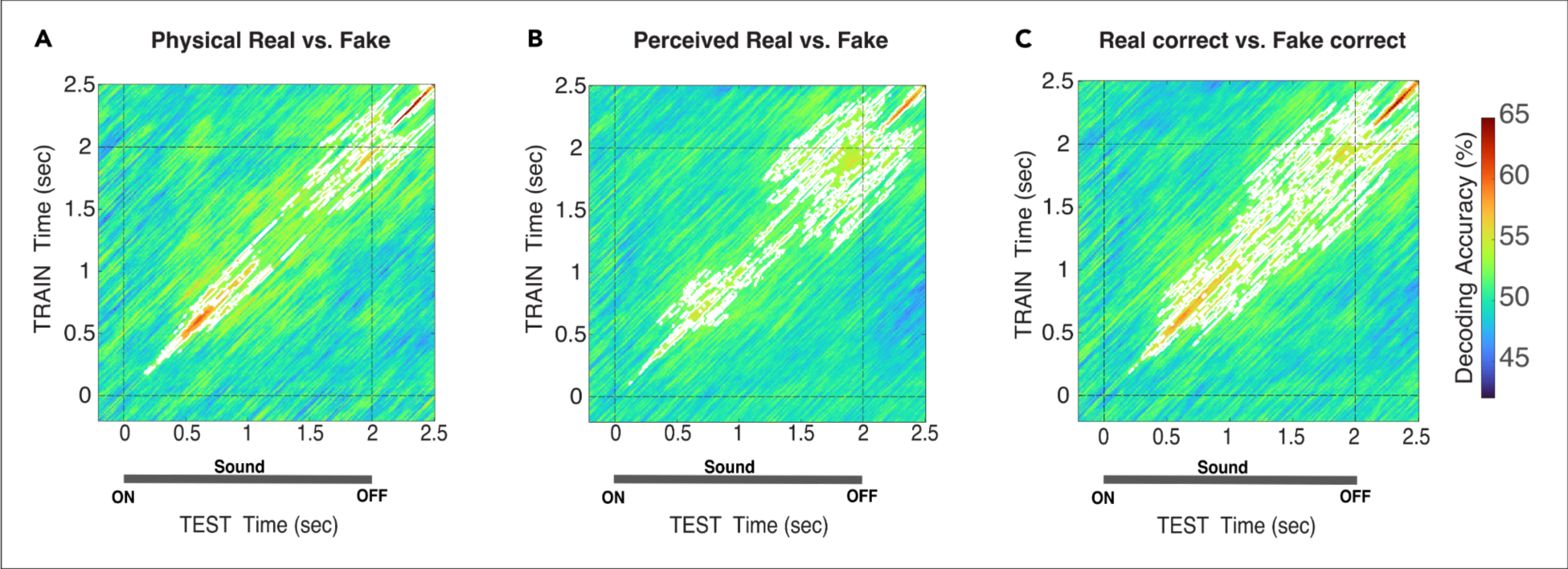
Temporal generalization matrix of *Real* vs. *Fake* decoding according to *physical stimuli, perceptual report*, and *correct response*. White contours indicate significant clusters across subjects and dashed lines at zero and two seconds indicate stimulus onset and offset. For all statistics, N = 20; *t-test* against 50%, cluster-definition threshold, *p* < 0.05; 1000 permutations.

### Spatial distribution of reverberant perceptual judgments

The decoding analyses reported in the above sections included all electrodes to maximize the available signal for analysis. Based on our whole-brain findings, we developed the working hypothesis that neural responses are dissociable into sensory (Binder et al., 2004; Desai et al., 2021; Lütkenhöner & Steinsträter, 1998; Näätänen et al., 1978) and higher-level perceptual decision-making phases (Diaz et al., 2017; Herding et al., 2019; Tagliabue et al., 2019), supported by distinct neural populations. Thus, we explored the spatial distribution of the processing cascade by examining two non-overlapping sensor clusters: temporal electrodes (pooled across hemispheres) and centroparietal electrodes (Figure 5).

**Figure 5:**
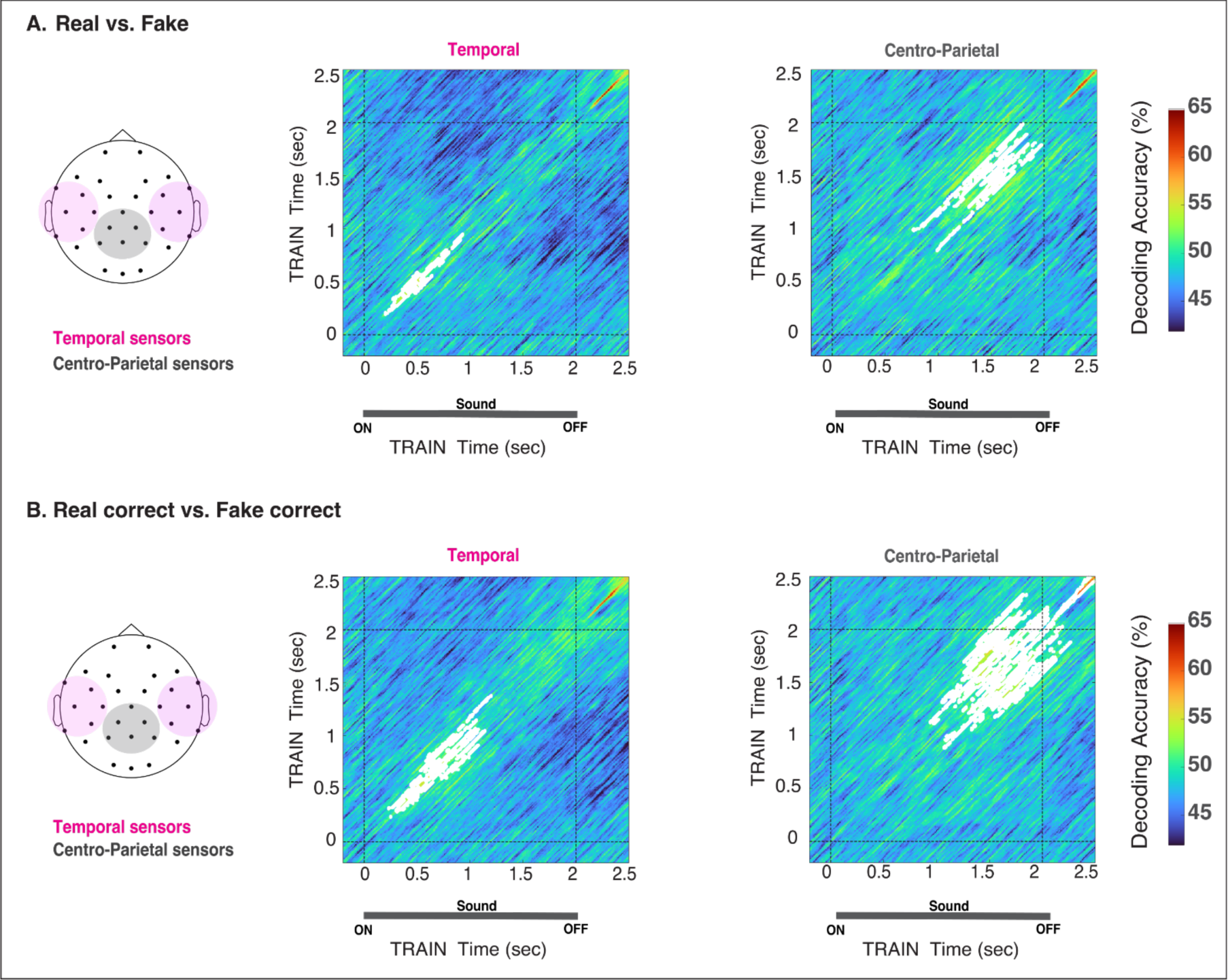
Temporal Generalization Matrices from two subsets of electrodes. The temporal cluster, pooled across electrodes and marked in pink in the layout, comprised the following electrodes: FC5, FC6, FT9, FT10, C3, C4, CP5, CP6, T7, T8, TP9, and TP10. The centro-parietal cluster, marked in gray in the layout, included Cz, Pz, P3, P4, CP1, and CP2. A: Real versus Fake (*Physical stimuli*). B: Real correct versus Fake correct (*Correct*). The white regions in the TGMs indicate significance. For all statistics, N = 20, *t-test* against 50%, cluster-definition threshold, *p* <0.05, 1000 permutations.

We performed temporal generalization analysis in these clusters to capture both temporal and representational dynamics of the Real vs. Fake decoding signal (Physical and Correct trial labels). As seen in Figure 5, the temporal clusters for both pairwise comparisons consistently showed significant decoding accuracy early in the trial (200–980 ms and 250–1400 ms), whereas the centro-parietal cluster showed significance later in the trial (850–1900 ms and 1100–2500 ms). For correct trials, significance was prolonged until the end of the epoch and beyond in the centro-parietal cluster, consistent with perceptual decision-making process indexed by the centro-parietal potential visible in these electrodes (Kelly & O’Connell, 2013; Tagliabue et al., 2019). These patterns suggest dissociable temporally and functionally specific representations in reverberant perceptual judgments.

## DISCUSSION

Reverberation is a ubiquitous acoustic feature of everyday environments, but its perception and neural representations remain understudied in comparison to other sounds. This study examined the dynamics of EEG responses to natural acoustic environments while human listeners classified real and synthetic reverberant IRs convolved with speech samples. Participants reliably categorized real and synthetic reverberations, which is consistent with prior work using similar stimuli (Traer & McDermott, 2016). In two temporal windows, starting at ∼500 ms and later around ∼2000 ms, neural response patterns reliably distinguished *Real* and *Synthetic* IRs (Figure 2B), with higher classification accuracy when subselecting trials for correct responses (Figure 3B). The two regimes of significant classification did not generalize to each other, indicating dissociable neural representations in the early and late phases of each trial (Figure 4). Moreover, the early and later components mapped to temporal and centro-parietal electrode clusters, respectively, suggesting distinct loci of underlying activity (Figure 5). Our findings demonstrate that at least two sequential and independent neural stages with distinctive neural operations underpin the perception of natural acoustic environments.

By definition, real-world spaces have real-world acoustics; people do not routinely discriminate between real and synthetic reverberation in everyday life. However, people do perceive and process reverberation, overtly or covertly, as an informative environmental signal rather than simply a nuisance distortion. Measuring authenticity perception by systematically manipulating statistical deviation from ecological acoustics probes the dimensionality and precision of the human auditory system’s world model (Traer & McDermott, 2016). Here, we extend previous work by tracking the neural operations that facilitate these judgments. Our EEG results suggest an ordered processing cascade in which sensory representations are integrated and abstracted into post-sensory or decisional variables over the course of reverberant listening. Tentatively, these results are consistent with the interpretation that stimulus acoustics influence, but do not directly drive, environmental authenticity judgments. Further research could extend the repertoire of acoustic statistics tested for perceptual sensitivity, including binaural cues (Młynarski & Jost, 2014; B. G. Shinn-Cunningham et al., 2005), as well as leverage imaging methods with greater spatial resolution to refine our picture of the reverberant perception circuit.

### Perceptual sensitivity to reverberant signal statistics

Our behavioral results replicate previous findings (Traer & McDermott, 2016), confirming that listeners are remarkably sensitive to statistical regularities of reverberation that allow them to distinguish real from synthetic acoustic environments. Human listeners reliably identified both *Real* and *Fake* trial types above chance, with higher accuracy for *Real* relative to *Fake* IRs. *Real* IRs have a more consistently identifiable statistical profile, whereas *Fake* variants could mimic that profile or deviate from it in specific ways. For example, *Ecological* IRs were intentionally crafted to emulate physical acoustics, while *Time-Reversed* were very saliently deviant and easily identified as *Fakes* (>95% accuracy). Thus, we would expect this pattern in the one-interval forced-choice (1IFC) paradigm we employed to accommodate the practical constraints of an EEG experiment. Traer et al. (Traer & McDermott, 2016) used a 2AFC task comparing real and synthetic IRs in the same trial, in which subjects may have selected “the most realistic IR” among the two stimuli. In contrast, in our task, participants heard only a single IR sample per trial and thus likely had to compare the stimulus with an internal representation (template) of natural environments. Further research could investigate the nature of such a template or model, e.g., whether it is an innate filter from the auditory system or learned and shaped through development and experience (Młynarski & McDermott, 2019; Woods & McDermott, 2018). In addition, while our monaural stimuli faithfully captured and reproduced the spectrotemporal statistics of real reverberation, they omitted the binaural spatial cues (e.g., interaural time, level, and interaural correlation differences) that carry important perceptual information (Młynarski & Jost, 2014; B. Shinn-Cunningham, 2005; B. G. Shinn-Cunningham et al., 2005) and may be investigated in future work.

### Dissociating sensory and perceptual decisional processing stages during reverberant authenticity judgments

Our analysis of the neurodynamics of natural environmental acoustics revealed at least two critical time windows that reliably distinguish real and synthetic IRs. Furthermore, the TGM geometric patterns (Fyshe, 2020; King & Dehaene, 2014) indicate two sequentially distinct neural operations with their own internal configuration and locus. The early decoding regime started shortly after stimulus onset, with a maximum peak of around 600 ms (Figure 2). Unlike the second decoding regime, it was invariant to subjective reports and response accuracy, as shown in Figures 3A–B. Consistent with previous research linking the encoding of low-level auditory signal features to the early stages of processing in the auditory cortex (Bidelman et al., 2018; Binder et al., 2004; Desai et al., 2021; Lütkenhöner & Steinsträter, 1998; Näätänen et al., 1978), the sensor space TGM showed that the early decoding phase, but not the later phase, was reliably localized to temporal sensor clusters (Figure 5). Furthermore, the timing of this regime aligns with previous research that identified modulatory effects on the neural response to reverberant sounds even when reverberation was unrelated to the task, underscoring the significance of this period for sensory-driven effects rather than task-driven effects (Bidelman et al., 2018; Puvvada et al., 2017; Teng et al., 2017). Taken together, these results suggest that the early decoding regime reflects early sensory processing of reverberant acoustic features. The second regime of significant decoding began in the peri-offset period, preceding the response cue. It was modulated by perceptual reports and response accuracy and independent of stimulus features orthogonal to IR (i.e., speaker gender), motor preparation, or mapping response location (see Figure 3 and Figure S3 in supplementary material). Notably, the TGMs shown in Figure 4 indicate that the pattern underlying the second regime is not generalized from the decoding earlier in the trial, indicating a distinct transformation of previous stimulus representations. The cascade of information processing is initially driven by sensory features, but perceptual representations could, for example, index discrete or probabilistic decision values, updating dynamically (Kelly & O’Connell, 2013; O’Connell et al., 2018; Romo & de Lafuente, 2013).

The modulation of decodability as a function of trial and response type—physical, correct, and incorrect—supports this notion, even though incorrect trials were not sufficient to allow for a fair comparison. Thus, the second decoding regime near stimulus offset likely reflects higher-level decision processes underlying the perceptual judgments, not a re-entrant sensory representation (King & Dehaene, 2014). The decodability pattern was reliably observed in the cluster of centro-parietal sensors (Figure 5), consistent with research linking the centro-parietal positivity (CPP) potential to sensory evidence accumulation and perceptual decision-making (Kelly & O’Connell, 2013; Philiastides et al., 2014; Philiastides & Sajda, 2006; Tagliabue et al., 2019). Together, these observations align with prior research on auditory perception, showing that sensory information is shaped by expectations (Traer et al., 2021), attention (Alain & Arnott, 2000; Sussman, 2017), sensory experience (Alain et al., 2014; Pantev et al., 1998), and contextual variables of the acoustic environment to form a coherent perceptual representation (Kuchibhotla & Bathellier, 2018; Lowe et al., 2022). Overall, our findings indicate that a cascade of evolving, dissociable neural representations mapped at different cortical loci underlies the perception and discrimination of acoustic environments.

Finally, understanding the neural and perceptual correlates of reverberant acoustics also has applied implications, such as the efficient generation of perceptually realistic virtual acoustics in simulation, gaming, and music and film production—domains in which "real" vs. "fake" judgments are indeed made routinely, albeit often implicitly (Girón et al., 2020; Helmholz et al., 2022; Neidhardt et al., 2022; B. Shinn-Cunningham, 2005; Traer et al., 2021). In orientation and mobility (O&M) settings for blind and low-vision individuals (for whom audition may be the sole sensory modality for distal perception) (Kolarik, Cirstea, et al., 2013; Kolarik, Pardhan, et al., 2013), rapid and accurate simulation of room reverberation can inform assistive technology or training interventions customized to specific environments.

## Acknowledgments

This work was supported by the E. Matilda Ziegler Foundation for the Blind (S.T., H.G.L.) and Smith-Kettlewell Eye Research Institute’s C.V. Starr Fellowship Fund (H.G.L.). We thank Audrey Wong-Kee-You, Yavin Alwis, and James Traer for their helpful feedback and assistance on early versions of this work.

## Author Contributions

Designed research H.G.L.; S.T. Performed research H.G.L.; S.T., Analyzed data: H.G.L.; S.T. Wrote the paper H.G.L., S.T.

## SUPPLEMENTARY MATERIAL

### 1. Temporal versus spectral synthetic variants

Figure S1 plots mean behavioral performance across subjects of the temporal (Linear Decay and Time Reversed) and spectral (Flat Spectral Dependence and Inverted Spectral Dependence) synthetic variants. Wilcoxon signed-rank tests revealed that these two averages differed significantly (Z = 3.92, p< 0.0001). Overall, temporal variants were categorized more accurately than spectral ones as “fake” signals.

**Figure S1.**
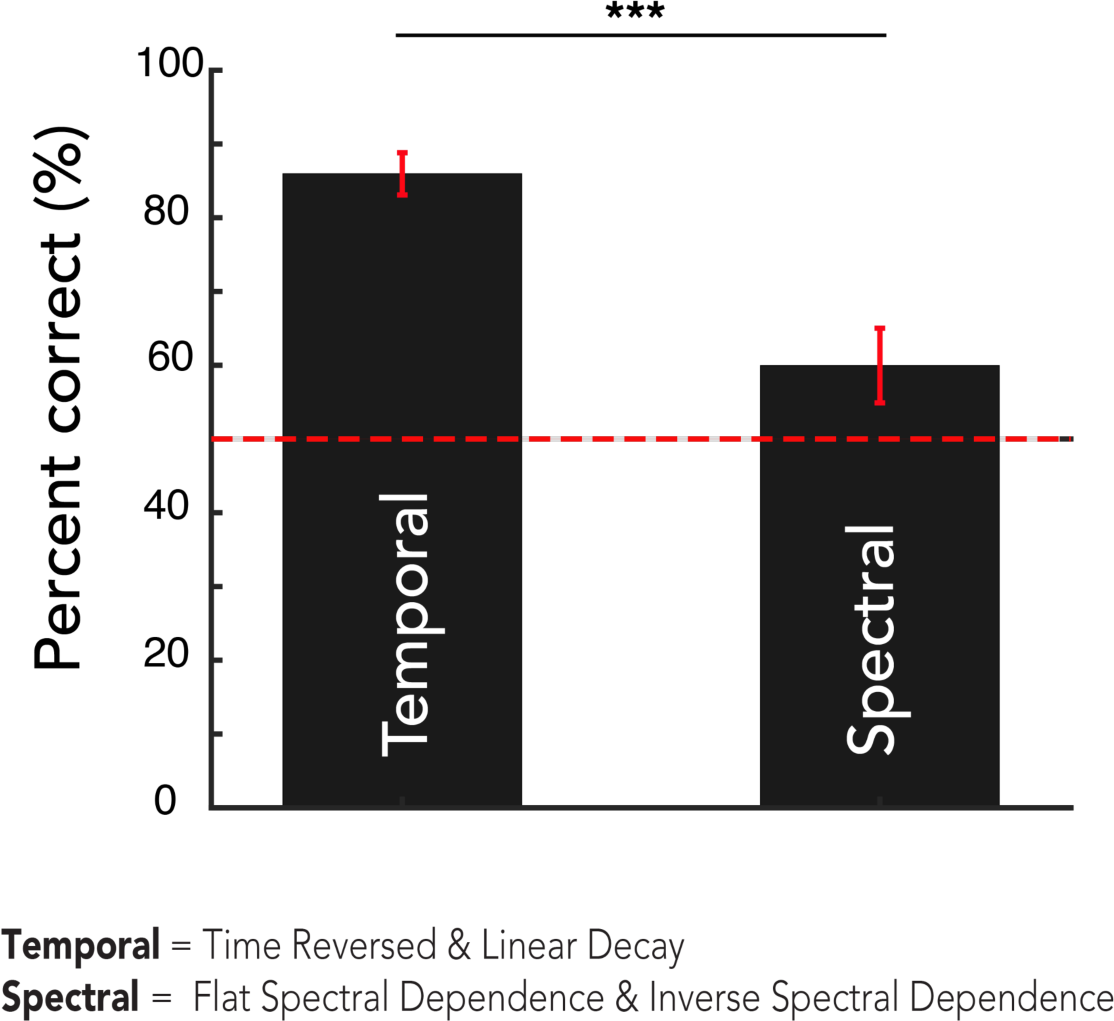
Group-level performance (mean accuracy ± s.e.m. across participants) averaged across temporal (Linear Decay and Time Reversed) and spectral (Flat Spectral Dependence and Inverted Spectral Dependence) fake variants. N = 20; non-parametric *signed-rank test*, threshold: \*\*\**p* < 0.001.

### 2. Real vs. Fake decoding time courses, excluding one at a time

Figure S2 shows the decoding time course for each pairwise comparison: *Real* vs. *Fake* variants except *Time-Reversed* (dark blue) reached significance from 530 ms to 780 ms and from 2140 ms to 2500 ms; *Real* vs. fake variants except *Linear Decay* (light blue) reached significance from 1460 ms to 1790 ms and from 2150 ms to 2470 ms; *Real* vs. fake variants except *Ecological* (green) reached significance from 510 ms to 970 ms and 2090 to 2500 ms; *Real* vs. fake variants except *Flat Spectral Dependence* (violet) became significant from 380 ms to 1100 ms and from 2200 ms to 2500 ms; and *Real* vs. fake variants except *Inverted Spectral Dependence* (pink) became significant from 3200 ms to 660 ms and from 2150 ms to 2500 ms.

**Figure S2.**
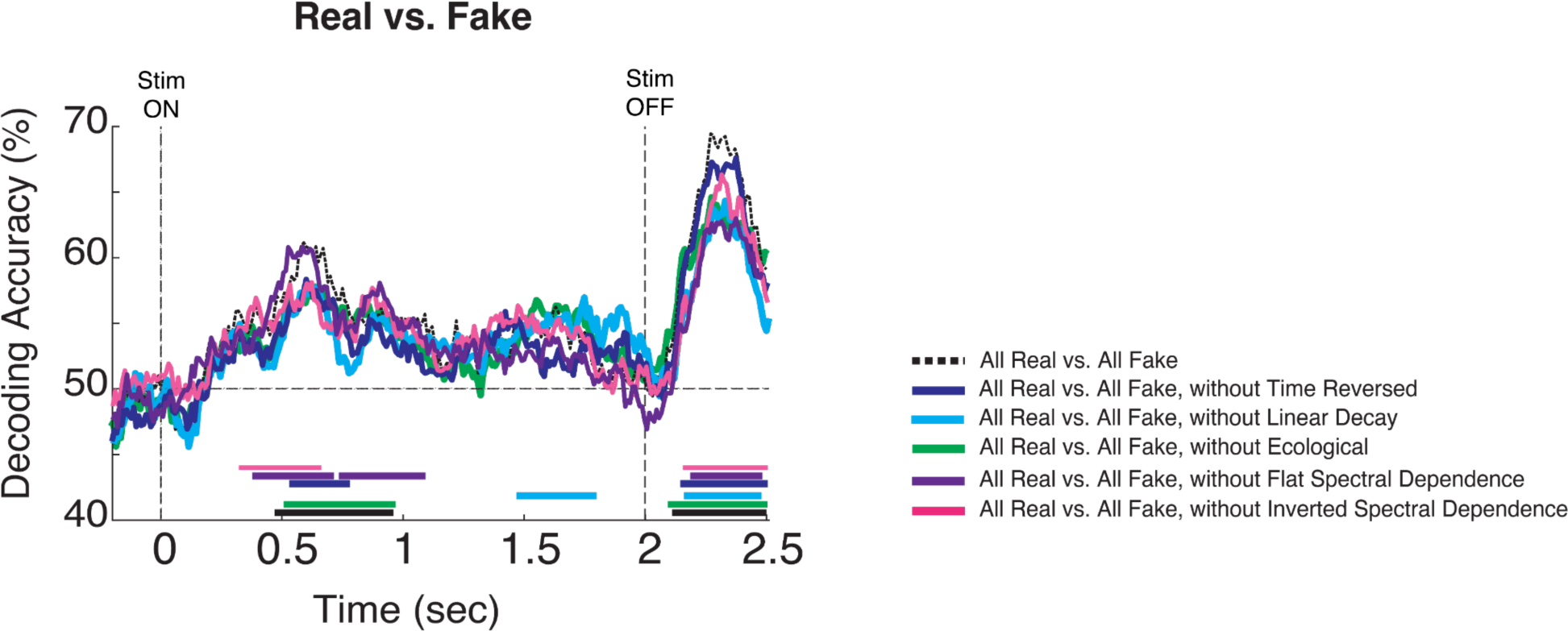
Decoding time courses of *Real* vs. *Fake* variants averaged across subjects. All *Real* vs. All *Fake* (black), All *Real* vs. All *Fake* without *Time Reversed* (dark blue), All *Real* vs. All *Fake* without *Linear Decay* (light blue), All *Real* vs. All *Fake* without *Ecological* (green), All *Real* vs. All *Fake* without *Flat Spectral Dependence* (violet), and All *Real* vs. All *Fake* without *Inverted Spectral Dependence* (pink). For all statistics, N = 20; *t-test* against 50%; cluster-definition threshold, *p* < 0.05; 1000 permutations.

### 3. Extraneous stimulus/response attribute and peri-offset decoding

Figures S3A and S3B show decoding time courses for trials labeled by impulse response type (room type) and speaker gender (male vs. female). No significant differences were found during the analyzed epoch. Figure S3C shows a decoding time course for trials labeled by response location. For this comparison, we extended the epoch of analysis up to 3.5 sec. Significance was reached from 2.7 to 3.5 secs.

**Figure S3.**
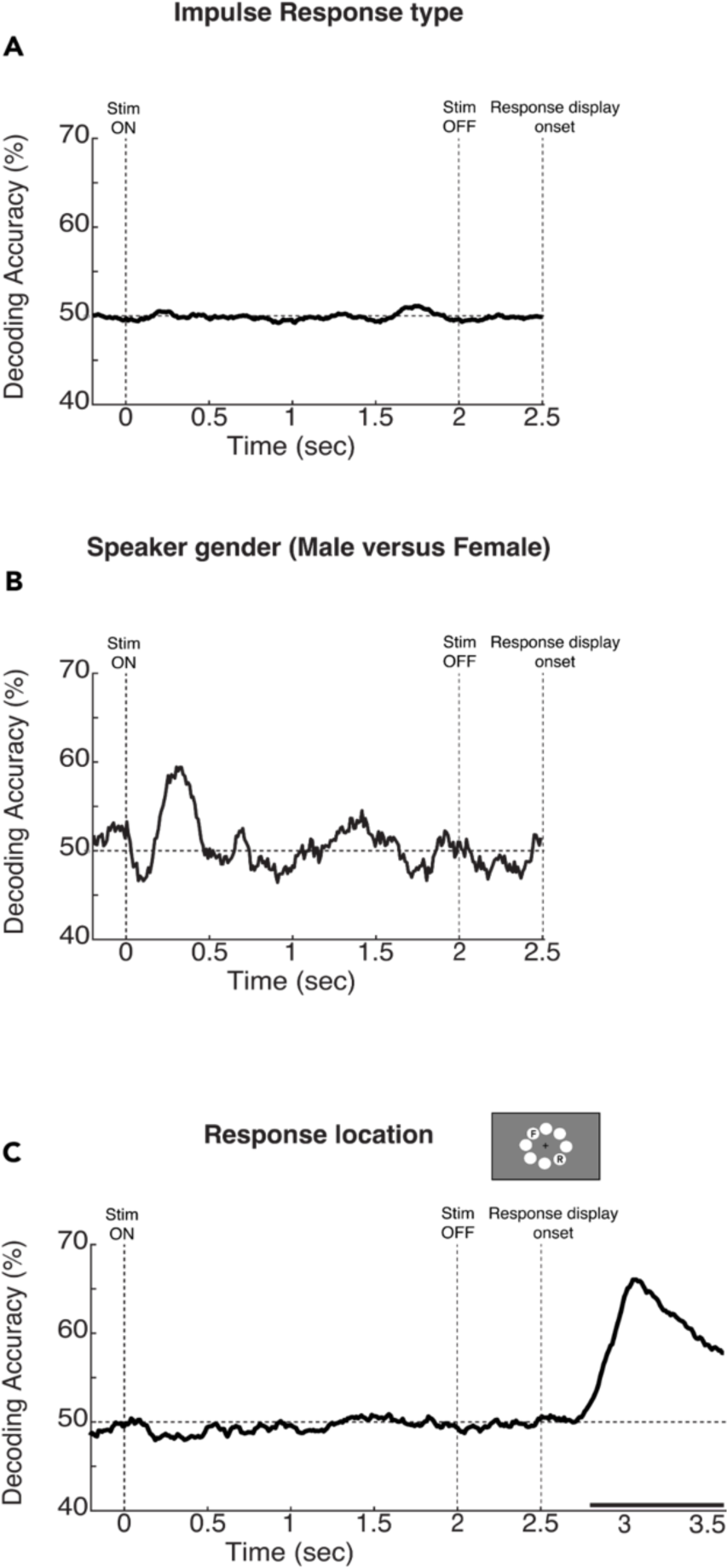
Decoding curves for IRs type, speaker gender (male versus female), and response location. A: Average of 435 pairwise comparisons between trials labeled with each of the 30 different IRs. Statistical analysis across subjects of the average of pairwise comparisons was not significant at any data point from 0 to 2500 ms. B: Decoding accuracy time courses for all trials labeled by speaker gender (Male versus Female). C: Decoding accuracy time courses for all trials labeled by response location. For all statistics, N = 20; *t-test* against 50%, cluster-definition threshold, *p* < 0.05; 1000 permutations.

